# Insights into the nature of the microalgal toxins from the *Chrysochromulina leadbeateri* blooms in Northern Norwegian fjords

**DOI:** 10.1101/2024.02.08.576530

**Authors:** Xinhui Wang, Mathias Fon, Aaron J.C. Andersen, Anita Solhaug, Richard A. Ingebrigtsen, Ingunn A. Samdal, Silvio Uhlig, Christopher O. Miles, Bente Edvardsen, Thomas O. Larsen

## Abstract

In May–June 2019, the microalga *Chrysochromulina leadbeateri* caused a massive fish-killing event in several fjords in Northern Norway, resulting in the largest direct impact ever on aquaculture in northern Europe due to toxic algae. Motivated by the fact that no algal toxins have previously been described from *C. leadbeateri*, we set out to investigate the chemical nature and toxicity of secondary metabolites in extracts of two strains (UIO 393, UIO 394) isolated from the 2019 bloom, as well as one older strain (UIO 035) isolated during a bloom in Northern Norway in 1991. Initial LC–DAD–MS/MS-based molecular networking analysis of the crude MeOH extracts of the cultivated strains showed that their profiles of small organic molecules, including a large number of known lipids, were very similar, suggesting that the same class of toxin(s) were likely the causative agents of the two harmful algal bloom (HAB) events. Next, bioassay-guided fractionation using the RTgill-W1 cell line and metabolomics analysis pointed to a major compound affording [M+H]^+^ ions at *m*/*z* 1399.8333 as a possible toxin, corresponding to a compound with the formula C_67_H_127_ClO_27_. Moreover, our study unveiled a series of minor analogues exhibiting distinct patterns of chlorination and sulfation, together defining a new family of compounds, which we propose to name leadbeaterins. Remarkably, these suspected toxins were detected *in situ* in samples collected during the 2019 bloom close to Tromsø, thereby substantiating their likely role in fish kills. The elemental compositions of the putative *C. leadbeateri* ichthyotoxins strongly indicate them to be long linear polyhydroxylated polyketides, structurally similar to sterolysins, reported from a number of dinoflagellates.

## 1. Introduction

Harmful algal blooms (HABs) pose a significant ecological and economic threat to the aquaculture industry, with recurrent episodes causing massive losses in the coastal waters of northern Europe, including the Baltic Sea, Kattegat–Skagerrak, eastern North Sea, Norwegian Sea, and the Barents Sea (Karlson et al., 2021). In most Scandinavian cases, these fish-killing events are attributed to marine haptophytes, and particularly to species belonging to the genera *Prymnesium* and *Chrysochromulina* (Johnsen et al., 2010; John et al., 2022). A historic HAB event in May–June 1988, caused by *Prymnesium polylepis* (formerly *Chrysochromulina polylepis*) devastated the Kattegat–Skagerrak and eastern North Sea ecosystems, leading to significant mortality in plankton, benthic organisms, and both wild and farmed fish (Sieburth and Johnson,1989; Edvardsen and Paasche, 1998; Gjøsæter et al., 2000; Cembella et al., 2005; Karlson et al., 2021). Initial documentation of *C. leadbeateri* as a causative species in the fish farm mortalities dates back to late May 1991, with incidents occurring within the Lofoten archipelago and Vestfjorden (Rey, 1991). In 2019, this species again formed a massive fish-killing bloom in early May–June in Vestfjorden in the Lofoten area (Vesterålen) and further north near Tromsø. The event resulted in the death of approximately 14,500 t of farmed salmon in Nordland and Troms counties, causing significant damage to the aquaculture sector (Karlsen et al., 2019; Marthinussen et al., 2020; Samdal and Edvardsen, 2020). Economic losses of the damage have been estimated to be up to 2.4 billion NOK, equivalent to approximately 220 million USD (Marthinussen et al., 2020). This is the largest fish kill caused by a HAB event ever recorded in Norway and, at least in economic terms, in northern Europe (Samdal and Edvardsen, 2020; John et al., 2022).

A common feature of many fish-killing microalgal species seems to be their capability to produce polyketide-derived lipophilic lytic compounds that damage gill tissues in fish (necrosis), resulting in increased ion permeability or suffocation, and death (Hallegraeff et al., 2023). The RTgill-W1 cell line is a well-characterized and relevant cell line derived from epithelial gill cells of the rainbow trout (*Oncorhynchus mykiss*), making it suitable for discovery of ichthyotoxins in bioassay-guided fractionation (Bols et al., 1994; Dorantes-Aranda et al., 2011). Despite progress in characterization of the chemical structures of the toxins of *Karlodinium parvum* (karlotoxins), *Prymnesium parvum* (prymnesins), and *Karenia brevisulcata* (brevisulcenals), the exact mechanisms by which these microalgae cause finfish mortality are not yet fully elucidated (Manning and La Claire, 2010; Van Wagoner et al., 2010; Deeds et al., 2015; Rasmussen et al., 2016; Binzer et al., 2019). However, the fact that the mode of action of karlotoxins has been suggested to be related to the disruption of the gill cell membrane by specific binding to, for instance, cholesterol, supports that they facilitate creation of pores in the membrane (Waters et al., 2015).

With no ichthyotoxins previously described from *C. leadbeateri*, the aim of the present study was to explore three strains of *C. leadbeateri* that have been isolated from algal blooms in 1991 and 2019 in order to: 1, obtain new insights into the chemical compositions of the strains using high-resolution mass spectrometry and molecular networking analysis, and; 2, employ the RTgill-W1 gill cell bioassay to tentatively identify ichthyotoxins produced by *C. leadbeateri*.

## 2. Materials and Methods

### 2.1 Reagents and standards

All solvents were purchased from Sigma-Aldrich and were HPLC grade for extraction work. For LC-HRMS, all solvents were LC-MS grade. Water was purified using a Milli-Q system (Millipore, Bedford, MA, USA). Leibovitz’s L-15 Glutamax medium, fetal bovine serum, penicillin/streptomycin, TrypLE, and phosphate buffered saline (PBS) were from Gibco (Thermo Fisher, Waltham, MA, USA).

### 2.2. Cultivation of *C. leadbeateri* cultures

*C. leadbeateri* UIO 035, 393 and 394 (**Table 1**) were obtained from the Norwegian Culture Collection of Algae (NORCCA, Oslo, Norway). The strains were cultivated in autoclaved f/2 medium of salinity 30 (Guillard and Ryther, 1962) at 15 °C and an irradiance of 250 μmol m^−2^ s^−1^ with a light–dark cycle of 14:10 h. The cultures were grown in 10-L Nalgene polycarbonate bottles with 5-L volumes for 2 wk, with gentle aeration.

**Table 1.**
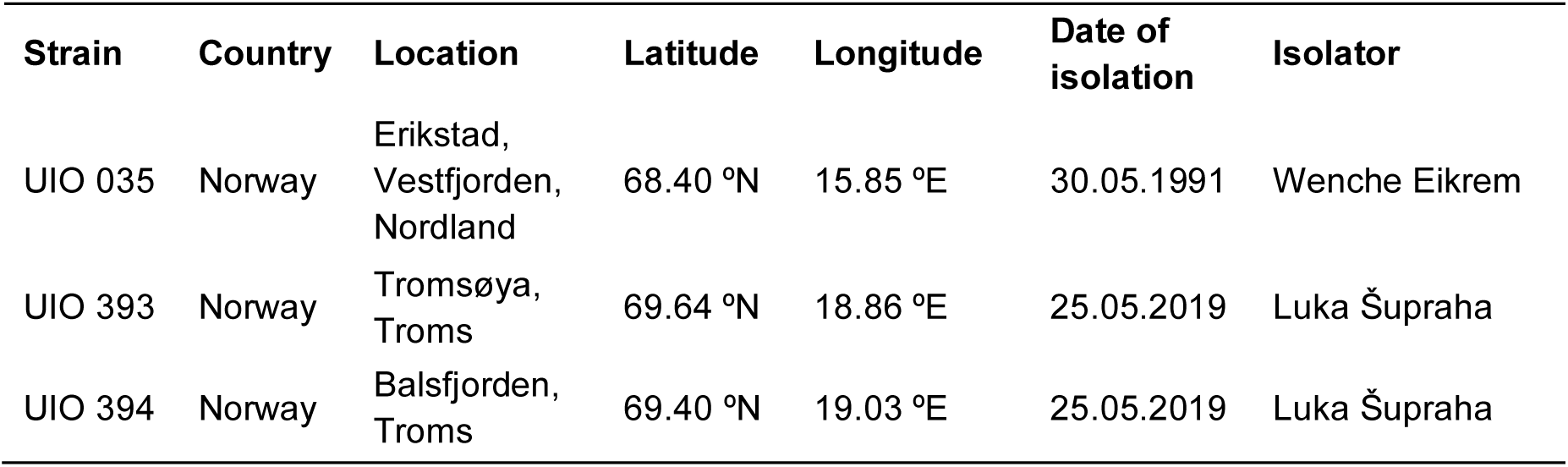
Details of sampling locations of *C. leadbeateri* isolates UIO 035, 393 and 394.

### 2.3. Harvest of biomass and extraction procedure

The cells were collected by gentle vacuum filtration using multiple glass microfiber filters (Whatman GF/C, 1.2 µm mesh size, diameter 47 mm; Sigma–Aldrich, St. Louis, USA), changed around every 300–400 mL depending on clogging. Biomass from the filters was extracted by adding 20 mL MeOH, briefly vortex-mixing, sonicating for 30 min, followed by centrifugation (15 min, 4220 *g*). The supernatant was decanted into polypropylene tubes, and the sediment was extracted a second time using the same procedure. The combined MeOH-extracts (40 mL) were evaporated to dryness under N_2_ at 35 °C.

### 2.4. Sample fractionation for RTgill-W1 cell assay

The MeOH extracts from cells of the three strains were subjected to two rounds of fractionation.

1. Initial fractionation of each extract was performed using an Isolera One automated flash system (Biotage, Uppsala, Sweden) with a 10 g RP C_18_ (15 μm, 100 Å; Grace) column (Biotage), using MeOH and water both contain 0.1% formic acid as eluents (15 mL/min).

Gradient stepwise elution with 10, 20, 30, 40, 50, 60, 70, 80, 90, and 100% MeOH (45 mL each) resulted in 10 fractions. Based on LC–MS (section 2.6), the 10 fractions were combined into 5 fractions for each strain: 20 and 40% MeOH (fraction A), 50 and 60% MeOH (fraction B), 70 and 80% MeOH (fraction C), 90% MeOH (fraction D), and 100% MeOH (fraction E). The five fractions (A–E) were evaporated under vacuum for RTgill-W1 cell bioassay.

2. After analysis, the remainder of fraction D from UIO 394 and UIO 035 were combined, evaporated to dryness under a stream of nitrogen, and the residue (0.6 mg) dissolved in 100 µL 90% MeOH. The combined fraction D was then fractionated on a Gemini C6-phenyl column (5 μm, 110 Å, 250 × 10 mm, Phenomenex, Torrance, CA, USA) held at 40 °C, using a linear gradient (4 mL min−1) from 10–100% MeCN in Milli-Q water (both containing 0.1% formic acid) over 30 min by on an Agilent Infinity 1290 HPLC–DAD (Agilent Technologies, Santa Clara, CA, USA). Eluent was monitored by UV absorbance at 230, 280, and 320 nm, and fractions were collected every 1 min to give 30 subfractions. To reduce the number of fractions for the RTgill-W1 cell bioassay, the fractions collected every two minutes from the initial set of 30 were combined to form 15 subfractions (fractions D1-D15).

### 2.5. RTgill-W1 cell assay

The epithelial gill cell line RTgill-W1 from rainbow trout (*Oncorhynchus mykiss*) (American Type Culture Collection (ATCC), Manassas, VA, USA) (Bols et al., 1994) was used to assess bioactivity in the crude extract and fractions from *C. leadbeateri*. The cell line was grown in Leibovitz’s L-15 Glutamax medium, supplemented with 10% fetal bovine serum) and 1% penicillin/streptomycin (Gibco, Thermo Fisher) in a non-ventilated 75 cm^2^ cell culture flask at 19 °C in a temperature-controlled incubator (IPP110+, Memmert, Germany). The cells were sub-cultured 1:2 once every 10 d using TrypLE. The cells were counted using a brightfield Cell Counter (DeNovix, Wilmington, DE, USA) and plated in 96-well plates (109,000 cells/cm^2^) 1— 2 d before the experiment. The cells were confluent by the day of exposure. The outer wells in the 96-well plate were not utilized for cell seeding but were filled with PBS (7.4 pH) to mitigate the edge effect (Mansoury et al., 2021).

Upon exposure, the medium was replaced with exposure-medium (L15ex with or without *C. leadbeateri* extract) containing 1% v/v of the fraction dissolved in MeOH. Additionally, medium was also replaced with L15ex medium containing 0.1% v/v of the fraction + 0.9% v/v MeOH (resulting in a total volume of 1% MeOH in the treatment). Appropriate controls containing the same percentage of MeOH were included in each experiment. In contrast to the L-15 growth medium, the L-15ex medium consists only of salts, pyruvate and galactose at similar concentrations to the L-15 medium, and the absence of amino acids often makes the assay more sensitive (Schirmer et al., 1997; OECD, 2021). Three technical replicates were performed and viability was measured after 24 h exposure using the Alamar Blue fluorometric assay according to the manufacturer’s protocol (Invitrogen, Thermo Fisher, Waltham, MA, USA). The dark blue oxidized form of Alamar Blue dye (resazurin) is reduced to a highly fluorescent product (resorufin), which is thus proportional to the number of viable cells. The fluorescent intensity (λ_ex_ 555 nm, λ_em_ 585 nm) was measured using a SpectraMax i3x plate reader (Molecular Devices, San Jose, CA, USA) after incubation for 2 h. Additionally, the morphology of the cells was visualized by light microscope (Zeiss Observer A1, Carl Zeiss, Oberkochen, Germany).

### 2.6. HPLC–DAD–HRMS

HPLC–DAD–HRMS analyses the MeOH extracts of cells, and all the fractions for RTgill-W1 cell assay, were performed after reconstitution in MeOH–water (90:10 v/v). The samples were transferred to 1.5 mL Eppendorf tubes and centrifuged (12,100 *g*, ambient temperature, 3 min) to remove residues of the glass fiber filters.

Separation was performed on an Ultimate 3000 UHPLC system (Dionex, Sunnyvale, CA, USA) including a DAD detector, using an InfinityLab Poroshell 120 Phenyl Hexyl (150 × 2.1 mm, 1.9 µm; Agilent Technologies, Santa Clara, CA, USA) at 40 °C and a flow rate of 0.4 mL min^−1^. A linear water–MeCN gradient, containing 20 mM formic acid, was applied from 10% to 100% MeCN over 10 min and maintained for 2 min before returning to the initial conditions. The samples were maintained at 8 °C in the autosampler, and 5 µL was injected for each analysis. Mass spectrometric data were acquired on a MaXis HD QTOF-MS equipped with an electrospray ionization source. The mass spectrometer was operated in positive mode with a capillary voltage of 3500 V, and data recorded in a scan range from *m*/*z* 100 to 1500 at a rate of 8 scans/s. The drying gas flow rate was set to 11.0 L min^−1^, the temperature to 220 °C and the nebulizer pressure was 2.0 bar. To accommodate larger ions, the following settings were used for the collision cell: transfer time 75 µs, collision cell RF 1100 Vpp, and prepulse storage 10 µs. The mass spectrometer was calibrated using sodium formate. Data were evaluated with Bruker Compass DataAnalysis Version 5.3 (Build 556.396.6383).

### 2.7. LC–DAD–QTOFMS analysis

Samples for the molecular networking analysis were analyzed using the Agilent LC–MS system. LC–DAD–QTOF-MS was performed on an Agilent Infinity 1290 UHPLC system (Agilent Technologies, Santa Clara, CA, USA) equipped with a diode array detector. Separation was achieved on a 150 × 2.1 mm, 1.9 μm, Poroshell 120 Phenyl Hexyl column (Agilent Technologies), held at 40 °C.

MS detection was performed on an Agilent 6545 QTOF-MS instrument equipped with an Agilent dual jet stream electrospray ion source (ESI) with a drying gas temperature of 250 °C in positive mode and 325 °C in negative mode, a gas flow of 8 L min^−1^, a sheath gas temperature and a flow rate at 300 °C and 12 L min^−1^, respectively. Capillary voltage was set to 4000 V and nozzle voltage to 500 V in both positive and negative modes. MS spectra were recorded as centroid data, scan range *m*/*z* of 100−1700, and auto MS/HRMS fragmentation was performed at three collision energies (10, 20, and 40 eV), on the three most intense precursor peaks per cycle. The acquisition rate was 10 spectra s^−1^. Data were handled using Agilent MassHunter Qualitative Analysis software (Agilent Technologies). Tributylamine (1 μM, Sigma–Aldrich) and hexakis(2,2,3,3-tetrafluoropropoxy)phosphazene (10 μM, Apollo Scientific Ltd., Cheshire, UK) in 70% MeOH were infused as lock mass compounds in the second sprayer using an extra LC pump at a flow rate of 15 μL min^−1^ using a 1:100 splitter, and the *m/z* of the protonated (*m*/*z* 186.2216 and 922.0098, respectively), or the *m/z* of [M + HCO_2_]^−^ of the phosphazene (*m*/*z* 966.0007) employed for internal mass calibration.

### 2.8. Molecular networking analysis

The MS/MS data files were converted from Agilent output (.d extension) to the mzXML file format with MS-Convert (https://proteowizard.sourceforge.io/download.html). The converted data files were uploaded to the GNPS Web platform (http://gnps.ucsd.edu) (Wang et al., 2016; Aron et al., 2020) for molecular network data analysis. The data was filtered by removing all MS/MS product-ions within ±17 *m*/*z* of the precursor *m*/*z*. MS/MS spectra were window-filtered by choosing only the top 6 product-ions in the ±50 *m*/*z* window throughout the spectrum. The precursor-ion mass tolerance was set to 0.05 *m*/*z* and the MS/MS product-ion tolerance to 0.10 *m*/*z*. A molecular network was then created where edges were filtered to have a cosine score above 0.68 and more than 4 matched peaks. Furthermore, edges between two nodes were kept in the network if and only if each of the nodes appeared in each other’s respective top 10 most similar nodes. Finally, the maximum size of a molecular family was set to 100, and the lowest scoring edges were removed from molecular families until the molecular family size was below this threshold. The spectra in the network were then searched against GNPS spectral libraries. The library spectra were filtered in the same manner as the input data. All matches kept between network spectra and library spectra were required to have a score above 0.68 and at least 4 matched peaks. The molecular networks were visualized using Cytoscape software (Shannon et al., 2003).

### 2.9. Data processing

The LC–MS data file was converted from Bruker output (.d extension) to the mzML file format with MS-Convert and subjected to MZmine 3 analysis (Schmid et al., 2023). The mass detection was performed using a centroid algorithm with a noise level of 400. The ADAP chromatogram builder was established using a minimum group size of 5 scans, a group intensity threshold of 400, a minimum highest intensity of 3000, and an *m*/*z* tolerance of 0.002. The peak deconvolution was performed using a local minimum feature resolver algorithm with the following standard settings: chromatographic threshold = 85%, minimum search range retention time = 0.02, minimum absolute height = 500, Min ratio of peak top/edge = 2, Peak duration range (min) = 0.00−1.20. Min number of data points = 5.

The chromatograms were deisotoped with an *m*/*z* tolerance of 0.001 and retention time tolerance of 0.02 min. The peak list was aligned using the join aligner algorithm with an *m*/*z* tolerance of 0.005, weight for *m*/*z* = 3, weight for retention time = 3, and retention time tolerance of 0.1 min. The peak list was eventually gap-filled using the peak finder module (intensity tolerance at 20%, *m*/*z* tolerance at 0.003, retention time tolerance at 0.02, and minimum data points at 3). Results were exported as a .csv file containing: row ID, *m*/*z*, RT and peak areas in each sample.

### 2.10. Statistical analysis

The data matrix was uploaded to the MetaboAnalyst 5.0 web site (http://www.metaboanalyst.ca) (Pang et al., 2022). The data set containing 3045 features was normalized by using Pareto scaling and submitted to principal component analysis (PCA). Total explained variance percentages of PC1 and PC2 were 27.4 and 20.2%, respectively. The dendrogram was obtained by hierarchical cluster analysis using the Euclidean distance and the “Ward” algorithm. One-way ANOVA was performed using a p-value <0.05 and post-hoc analysis, which determined the top 203 features with the most significantly variable mean for peak areas among the fractions. Hierarchical clustering analysis (HCA) of the top 100 features by using a heatmap was obtained using the Euclidean distance measure and the “Ward” algorithm for clustering. The chromatographic features were evaluated as areas under the curve (AUC) to represent feature abundance.

### 2.11. Dose–response modeling

The effects of crude *C. leadbeateri* extracts on cellular viability were estimated using a non-linear four-parameter log-logistic function. The function is defined as:

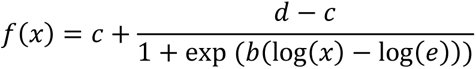

In this equation, f(x) represents the cellular viability expressed as percentage relative to control, and x denotes the concentration of algal cells, normalized against the sample. Parameter b describes the slope of the curve and e is the concentration at which the inflection point is found between c and d. The parameters c and d are constrained to 0 and 100, respectively. This constraint models the cellular viability within a range from 0% to 100% relative to the control. The *drc* package (Ritz et al., 2015) in R, a tool for dose–response analyses, was employed to fit this model to the data obtained from all independent replicates. From this model, the EC_50_ value, representing the concentration of algal cells at which the cellular viability is reduced to 50%, was estimated and serves as an indicator of the crude extract toxicity.

### 2.12. *In-situ* sample collection and preparation

A 900-mL seawater sample was collected from the shore of Hamna Harbour (GPS location: 69°41′58.5″N 18°58′10.4″E) and subsequently strained through a 100-µm mesh filter for biomass extraction. A 2-g Sep-Pak Vac C_18_ solid-phase extraction (SPE) column (Waters, Milford, MA, USA) was used, preconditioned with 10 mL of MeOH, and then with 10 mL of Milli-Q water. The filtered seawater (900 mL) was loaded onto the SPE column, the column was washed with Milli-Q water (10 mL), and the analytes were eluted with MeOH (15 mL).

The sample was evaporated in a ScanSpeed 40 vacuum centrifuge (LabogeneApS, Denmark) and the residue dissolved in MeOH (1 mL) and vortex-mixed for 10 s. The sample was transferred to a 2-mL Eppendorf tube, centrifuged (5 min, 13148 *g*), and the supernatant was transferred to a 1.5-mL HPLC vial for HPLC–DAD–HRMS analysis.

## 3. Results

In this study, we investigated MeOH extracts from cells of three cultured *C. leadbeateri* strains (UIO 035, UIO 393, UIO 394). These strains were isolated from algal blooms in Northern Norway in 1991 and 2019. Our aim was to compare their chemical profiles and identify potential toxins.

### 3.1. Chemical profiling of *C. leadbeateri* strains

Analysis of the MeOH extracts of the *C. leadbeateri* strains by LCMS/MS in positive ionization mode (section 2.7) revealed the chemical profiles of the three *C. leadbeateri* strains (**Figure S1**). Molecular networking elucidated the primary classes of molecules produced by these previously unexplored strains. The obtained molecular network consisted of 364 nodes, of which 30 were dereplicated against the GNPS library (**Figure 1**). Matching these nodes with the GNPS library allowed for the annotation of several nodes within the main network clusters (specifically cluster 1–3), including fatty amides, fatty acids, phospholipids (PCs), 1,2-diacylglyceryl-3-O-carboxy-(hydroxymethyl)-choline (DGCC), diacylglyceryl-trimethylhomoserine (DGTS) and other lipids. These analyses revealed the predominant constituents in *C. leadbeateri*. The dereplicated compounds are listed in **Table 2**.

**Figure 1.**
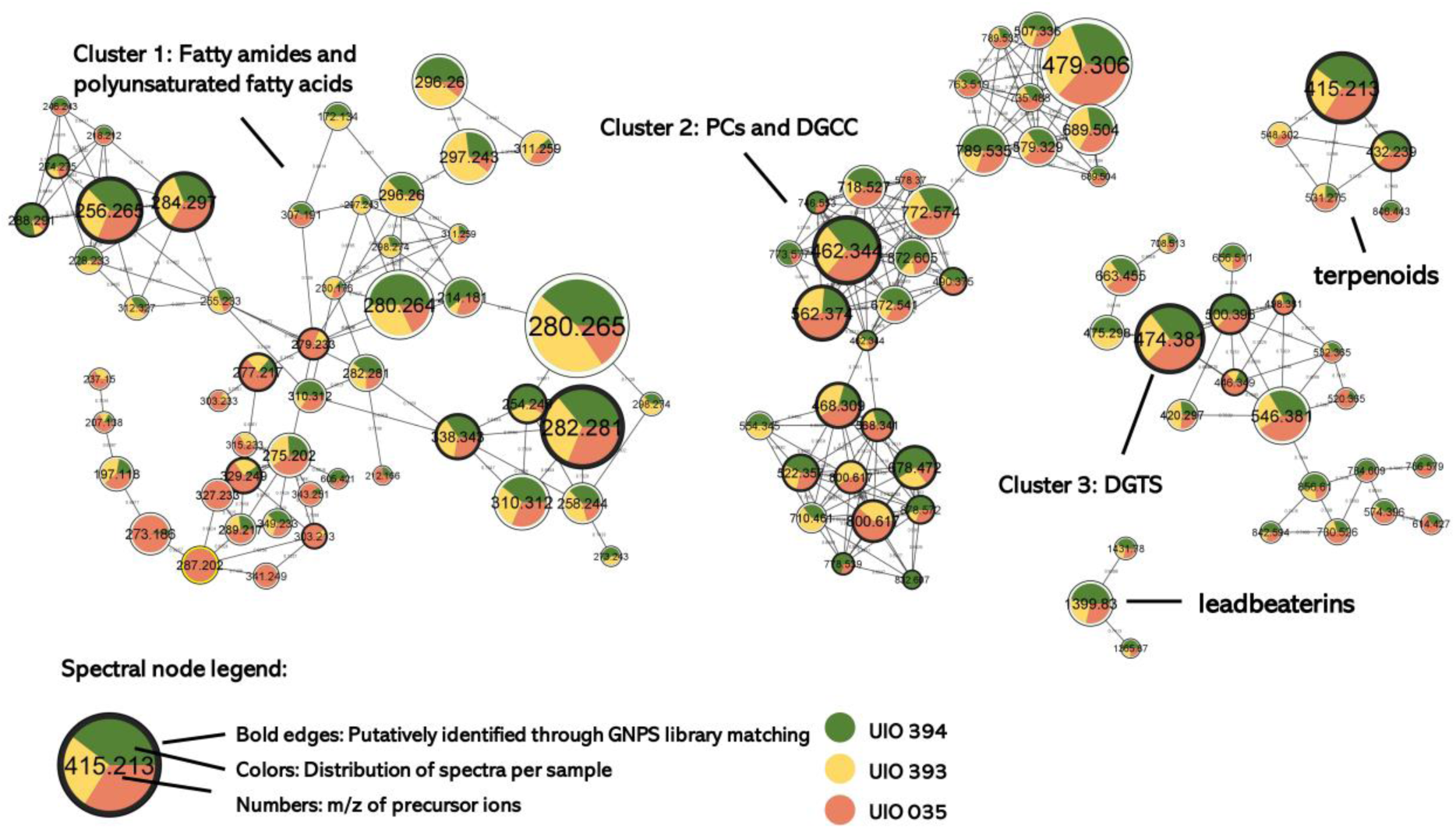
Molecular network of the MeOH extracts of biomass from cultures of the three *C. leadbeateri* strains. The nodes in the network are labeled with their respective precursor-ion *m*/*z* values in positive mode. The node size reflects the summed number of spectra in three strains.

**Table 2.** Overview of compounds tentatively identified in *C. leadbeateri* strains using the GNPS library.

| Compound class | Compounds name | Molecular Formula* | Ion | Accurate *m/z* | $\Delta$ (ppm) |
| --- | --- | --- | --- | --- | --- |
| Fatty amide | Octadecanamide | C<sub>18</sub>H<sub>37</sub>NO | [M + H]<sup>+</sup> | 284.2954 | 2.1 |
| Fatty amide | 9Z-Octadecenamide | C<sub>18</sub>H<sub>35</sub>NO | [M + H]<sup>+</sup> | 282.2795 | 1.4 |
| Fatty acid | Stearidonic acid | C<sub>18</sub>H<sub>28</sub>O<sub>2</sub> | [M + H]<sup>+</sup> | 277.2168 | 2.2 |
| Fatty acid | 9(10)-EpOME | C<sub>18</sub>H<sub>32</sub>O<sub>3</sub> | [M + H - H<sub>2</sub>O]<sup>+</sup> | 279.2318 | 0.0 |
| Fatty amide | Palmitamide | C<sub>16</sub>H<sub>33</sub>NO | [M + H]<sup>+</sup> | 256.2643 | 3.5 |
| Fatty acid | linoleic acid | C<sub>18</sub>H<sub>32</sub>O<sub>2</sub> | [M + Na]<sup>+</sup> | 303.2300 | 2.0 |
| Fatty acid | 13Z-Docosenamide | C<sub>22</sub>H<sub>43</sub>NO | [M + H]<sup>+</sup> | 388.3422 | 1.5 |
| Fatty amide | Lauryldiethanolamine\|2-[dodecyl(2-hydroxyethyl)amino]ethanol | C<sub>16</sub>H<sub>35</sub>NO<sub>2</sub> | [M + H]<sup>+</sup> | 274.2747 | 2.6 |
| Fatty acid | Docosahexaenoic acid | C<sub>22</sub>H<sub>32</sub>O<sub>2</sub> | [M + H]<sup>+</sup> | 329.2478 | 0.9 |
| Phospholipid | 1-Myristoyl-sn-glycero-3-phosphocholine | C<sub>22</sub>H<sub>46</sub>NO<sub>7</sub>P | [M + H]<sup>+</sup> | 468.3091 | 1.5 |
| Phospholipid | 1-(9Z-Octadecenoyl)-sn-glycero-3-phosphocholine | C<sub>26</sub>H<sub>52</sub>NO<sub>7</sub>P | [M + H]<sup>+</sup> | 522.3557 | 0.6 |
| Phospholipid | 1,2-Ditetradecanoyl-sn-glycero-3-phosphocholine | C<sub>36</sub>H<sub>72</sub>NO<sub>8</sub>P | [M + H]<sup>+</sup> | 678.5070 | 0.3 |
| Phospholipid | PC(22:6/22:6) | C<sub>52</sub>H<sub>80</sub>NO<sub>8</sub>P | [M + H]<sup>+</sup> | 878.5710 | 1.8 |
| Phospholipid | PC(18:1/19:1) | C<sub>45</sub>H<sub>86</sub>NO<sub>8</sub>P | [M + H]<sup>+</sup> | 800.6165 | 0.2 |
| Phospholipid | PC(14:0/22:6) | C<sub>44</sub>H<sub>76</sub>NO<sub>8</sub>P | [M + H]<sup>+</sup> | 778.5390 | 1.2 |
| Phospholipid | Lyso PC (22:6) | C<sub>30</sub>H<sub>50</sub>NO<sub>7</sub>P | [M + H]<sup>+</sup> | 568.3411 | 2.5 |
| Glycerolipid | Lyso DGCC (22:6) | C<sub>32</sub>H<sub>51</sub>NO<sub>7</sub> | [M + H]<sup>+</sup> | 562.3740 | 0.4 |
| Glycerolipid | Lyso DGCC (16:0) | C<sub>26</sub>H<sub>51</sub>NO<sub>7</sub> | [M + H]<sup>+</sup> | 490.3746 | 1.6 |
| Glycerolipid | Lyso DGCC (14:0) | C<sub>24</sub>H<sub>47</sub>NO<sub>7</sub> | [M + H]<sup>+</sup> | 462.3432 | 1.5 |
| Glycerolipid | DGCC (34:5) | C<sub>44</sub>H<sub>75</sub>NO<sub>8</sub> | [M + H]<sup>+</sup> | 746.5572 | 0.9 |
| Lipid | lysoDGTS 18:1 | C<sub>28</sub>H<sub>53</sub>NO<sub>6</sub> | [M + H]<sup>+</sup> | 500.3960 | 3.0 |
| Lipid | lysoDGTS 14:0 | C<sub>24</sub>H<sub>47</sub>NO<sub>6</sub> | [M + H]<sup>+</sup> | 446.3480 | 0.9 |
| Lipid | DGTS 18:2 | C<sub>28</sub>H<sub>51</sub>NO<sub>6</sub> | [M + H]<sup>+</sup> | 498.3798 | 1.8 |
| Lipid | DGTS 16:0 | C<sub>26</sub>H<sub>51</sub>NO<sub>6</sub> | [M + H]<sup>+</sup> | 474.3798 | 1.9 |
| Lipid | Commendamide | C<sub>18</sub>H<sub>35</sub>NO<sub>4</sub> | [M + H]<sup>+</sup> | 330.2641 | 0.9 |
| Piperidine derivative | 4-Hydroxy-1-(2-hydroxyethyl)-<br>2,2,6,6-tetramethylpiperidine | C<sub>11</sub>H<sub>23</sub>NO<sub>2</sub> | [M + H]<sup>+</sup> | 202.1803 | 1.0 |
| Terpenoid | 5-(1,2,4a,5-tetramethyl-7-oxo-3,4,8,8a-tetrahydro-2H-naphthalen-1-yl)-3-methylpentanoic acid | C<sub>20</sub>H<sub>32</sub>O<sub>3</sub> | [M + H]<sup>+</sup> | 321.2416 | -2.5 |
| Lipid | Monomyristin | C<sub>17</sub>H<sub>34</sub>O<sub>4</sub> | [M + H]<sup>+</sup> | 303.2530 | 0.3 |
| Phospholipid | 1-Myristoyl-2-hydroxy-sn-glycero-3-phosphoethanolamine | C<sub>19</sub>H<sub>40</sub>NO<sub>7</sub>P | [M + H]<sup>+</sup> | 426.2620 | 1.2 |
| Lipid | Dimethylsulfoniopropionate<br>(DMSP) | C<sub>5</sub>H<sub>10</sub>O<sub>2</sub>S | [M + H]<sup>+</sup> | 135.0473 | -0.7 |

\*Neutral formula

### 3.2. Bioassay-guided fractionation reveals potential toxins in *C. leadbeateri*

To assess the effect of the crude extract, an extract of *C. leadbeateri* strain UIO 394 was selected and prepared for initial toxicity screening on the RTgill-W1 cell line. Preliminary toxicity experiments showed similar results for strains UIO 035 and UIO 393. **Figure 2** illustrates the dose–response curve, showing a concentration-dependent reduction in cellular viability to RTgill-W1 cells after exposure to the crude extract for 24 h. The EC_50_ estimated from the four-parameter logistic model was 2.94 × 10^6^ cells mL^−1^ (CI 95%: ± 0.23 × 10^6^ cells mL^−1^).

**Figure 2.**
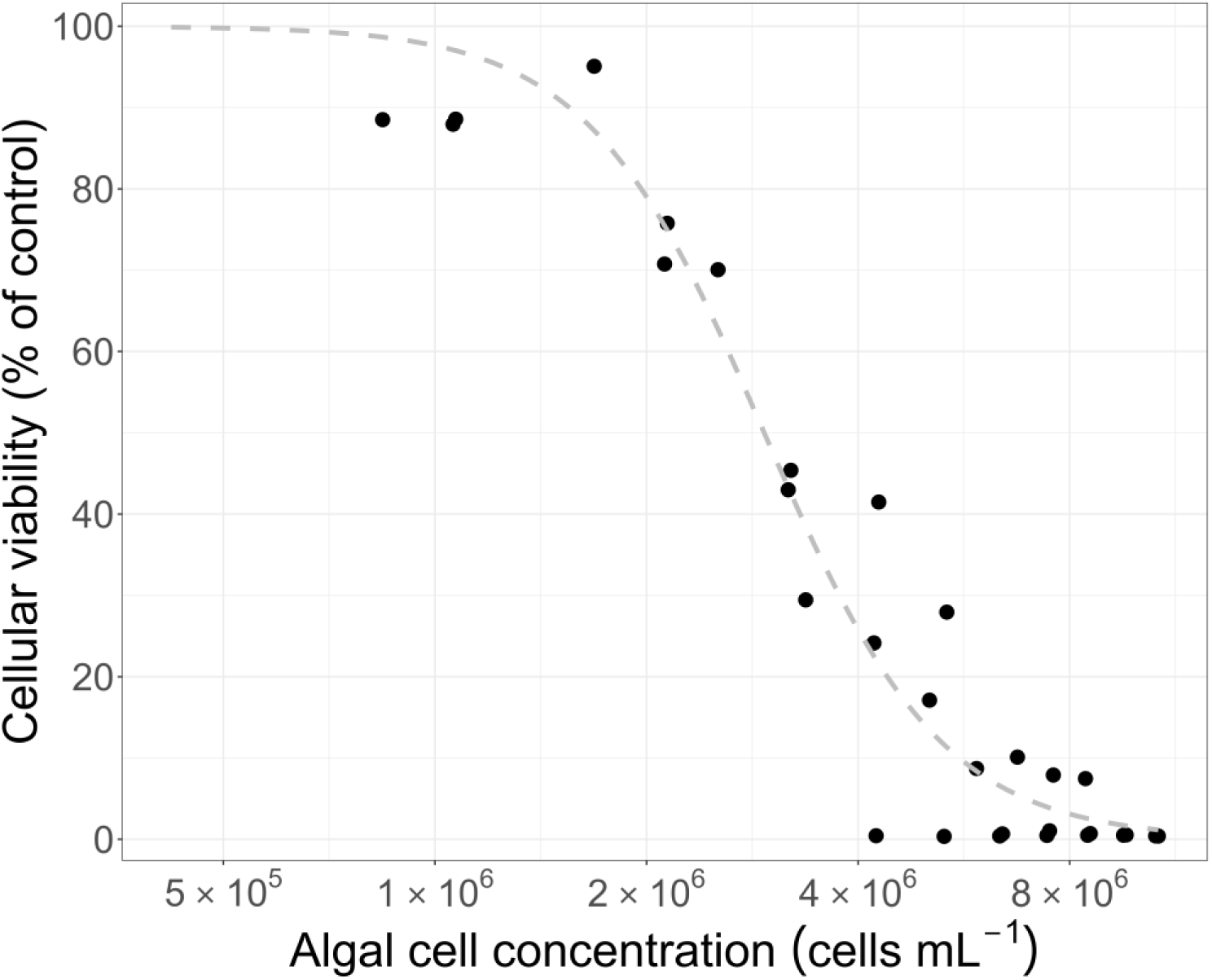
The cellular viability of RTgill-W1 cell line after 24 h exposure to crude extract from *C. leadbeateri* UIO 394. The exposure concentration was normalized to the algal cell concentration in the sample. The data was obtained from three independent experiments.

Next, we implemented a bioassay-guided fractionation approach, employing the RTgill-W1 gill cell line to screen for toxic compounds. The MeOH extracts from small-scale cultivation of each *C. leadbeateri* strain were subjected to fractionation, resulting in five fractions (A–E) per strain for bio-assay (**Table S1**). Concurrently, these fractions were subjected to analysis by LC–MS.

Bioassay results indicated the presence of highly toxic compounds in fraction D (eluted with 90% MeOH) in all strains (**Figure 3**). There was high activity at concentrations of 1% and 0.1% of fraction D. Additionally, high activity was observed at 1%, but not at 0.1% dilutions of fraction E. Further investigation of the LC–MS data revealed that bioactive fraction D of all three strains of *C. leadbeateri* exhibited strikingly similar chemical profiles (**Figure S2**). PCA (**Figure 4**) and a heatmap (**Figure 5**) generated using MetaboAnalyst 5.0 (Pang et al., 2022) reveal the potencial toxin within these bioactive fractions. Bioactive fraction D from all three strains grouped together, containing unique shared features that differed from those observed in the other fractions (**Figure 4**). In line with this, the heatmap analysis revealed specific compounds present in high abundance in fraction D (**Figure 5**). Comparison of the raw LC–MS data for the toxic and non-toxic fractions revealed an abundant [M+H]^+^ ion with *m*/z 1399.8333 in fraction D as a potential toxin. We propose to name this putative toxin leadbeaterin-1(**1**).

**Figure 3.**
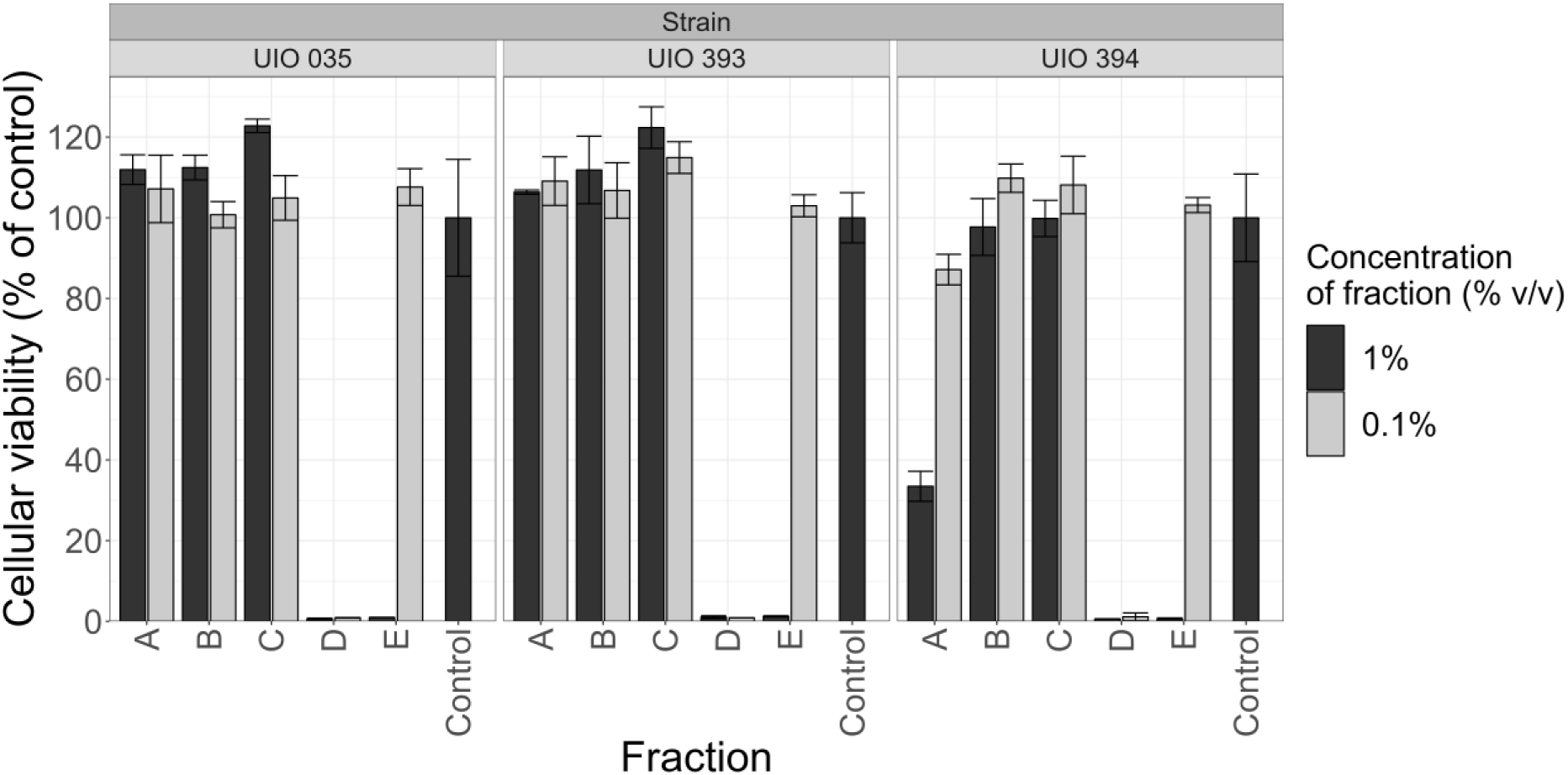
Analysis of the effects of fractions A–E on cellular viability of RTgill-W1 gill cells for three *C. leadbeateri* strains.

**Figure 4.**
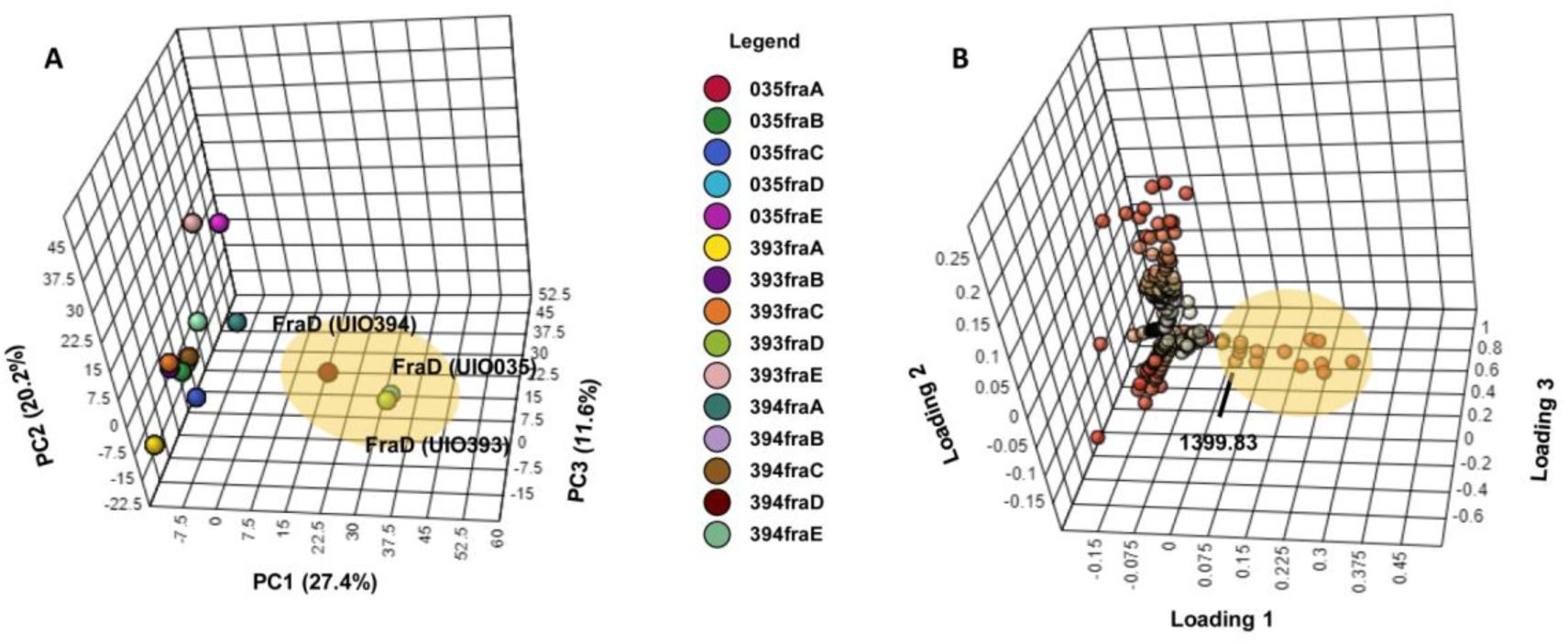
Principal component analysis of fractions from each *C. leadbeateri* strain. **A**. 3D scores plot. This plot visualizes the variance among the fractions from each strain along three principal components (PC1, PC2, and PC3). **B**. Loadings plot, highlighting the key metabolites that contribute to the observed variance among the fractions.

**Figure 5.**
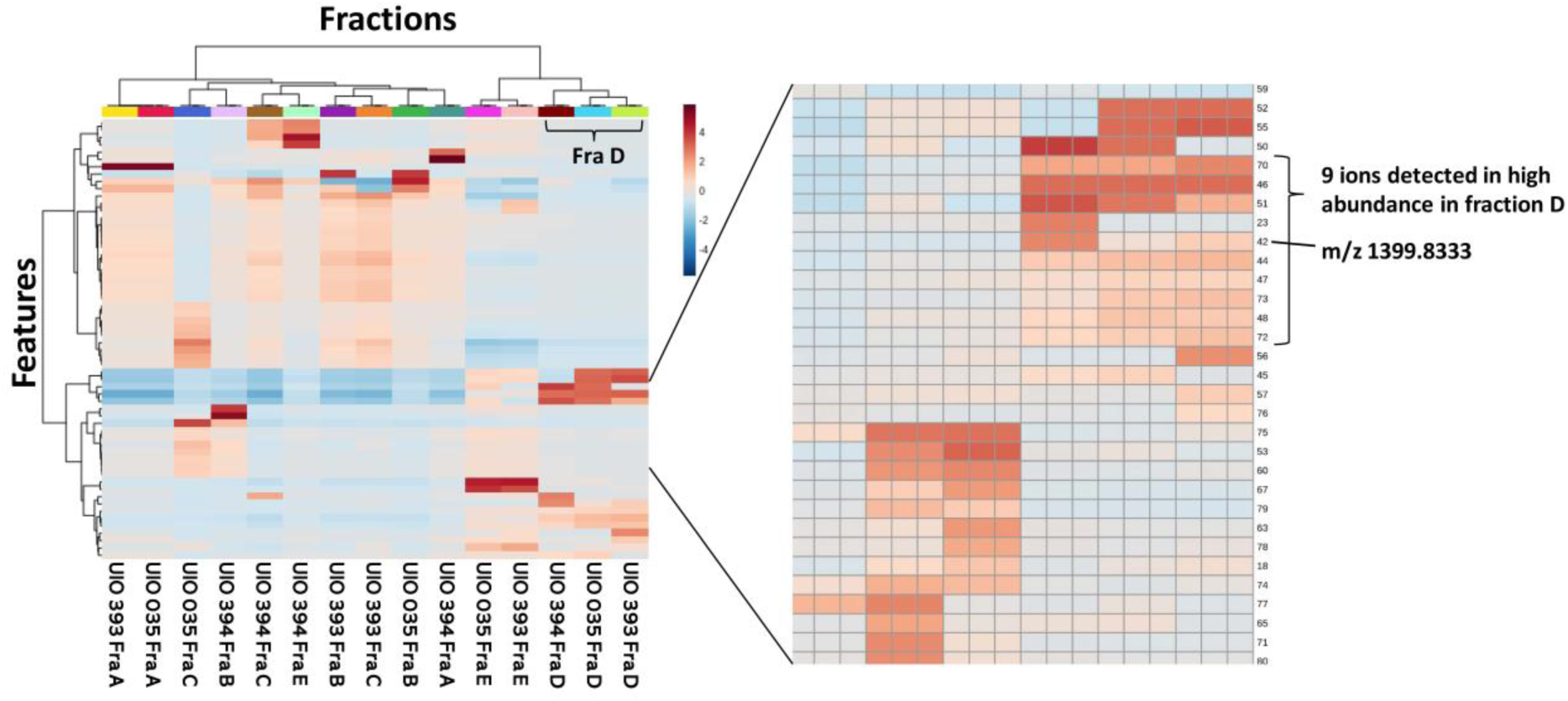
Hierarchical clustering analysis (HCA) of the 100 most significant features across the strains and fractions, visualized in a heatmap. The colors in the heatmap range from red, indicating high AUC in the original HPLC chromatograms, to blue, indicating low AUC value. The heatmap was generated using Pareto scaling for the AUC values of the top 100 features. “Fra D” denotes features for the highly toxic fraction D.

The bioactive fraction D combined from the UIO 394 and UIO 035 strains was subjected to further fractionation by semi-preparative HPLC resulting in 15 subfractions (fractions D1–D15). When the subfractions were tested with RTgill-W1 gill cells, only fraction D7 exhibited high bioactivity (**Figure 6**), suggesting that it contained the toxins. Furthermore, the LC–MS analysis revealed that more than 99% of the leadbeaterin-1(**1**) was present in toxic fraction D7(**Figure S3**), with the remainder detected in the weakly toxic fraction D8. No leadbeaterin-1(**1**) was detected in any of the other fractions. These results therefore suggested that leadbeaterin-1(**1**) was responsible for the bioactivity.

**Figure 6.**
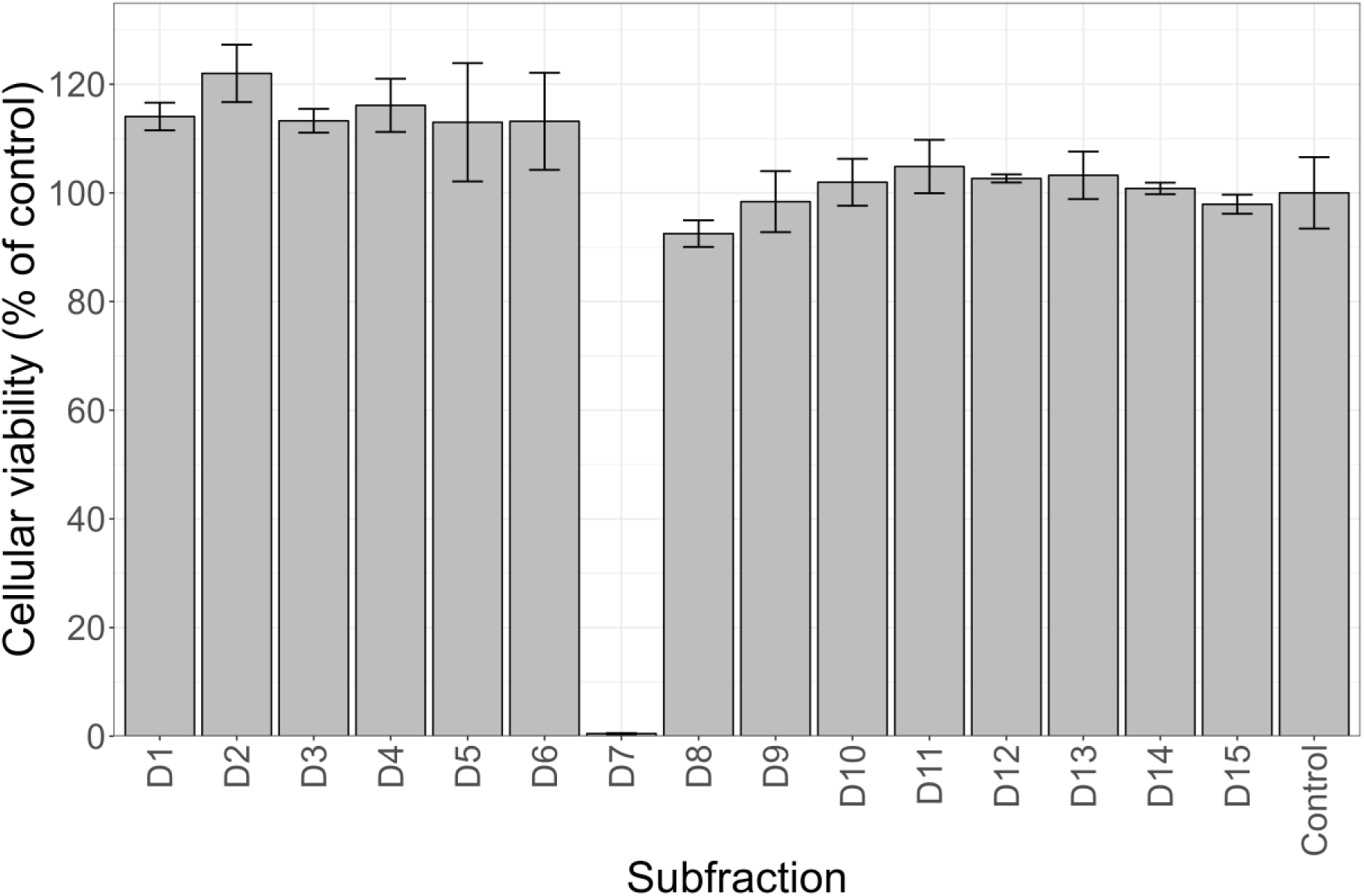
Cellular viability of RTgill-W1 gill cells treated with fractions D1–D15.

### 3.3. LC–MS/MS detection of potential toxin reveals new variants

With the evidence suggesting that the leadbeaterin-1(**1**) was the most likely candidate ichthyotoxin, a detailed analysis of the LC–MS data was undertaken to establish the elemental composition of the toxin based on accurate mass measurements, including the isotopologue patterns of the different adduct ions, i.e., [M + H]^+^, [M + Na]^+^, [M − H]^−^, and [M − 2H + Na]^−^ in positive and negative ionization mode (**Figure 7**). Only one plausible candidate emerged for both ionization modes with the NRC molecular formula calculator (https://metrology.shinyapps.io/molecular-formula-calculator/, v.1.01). The remaining candidates showed poor match to the experimentally observed HRMS spectra for both [M + H]^+^ and [M − H]^−^, thus establishing that the only acceptable elemental composition for leadbeaterin-1(**1**) is C_67_H_127_ClO_27_.

**Figure 7.**
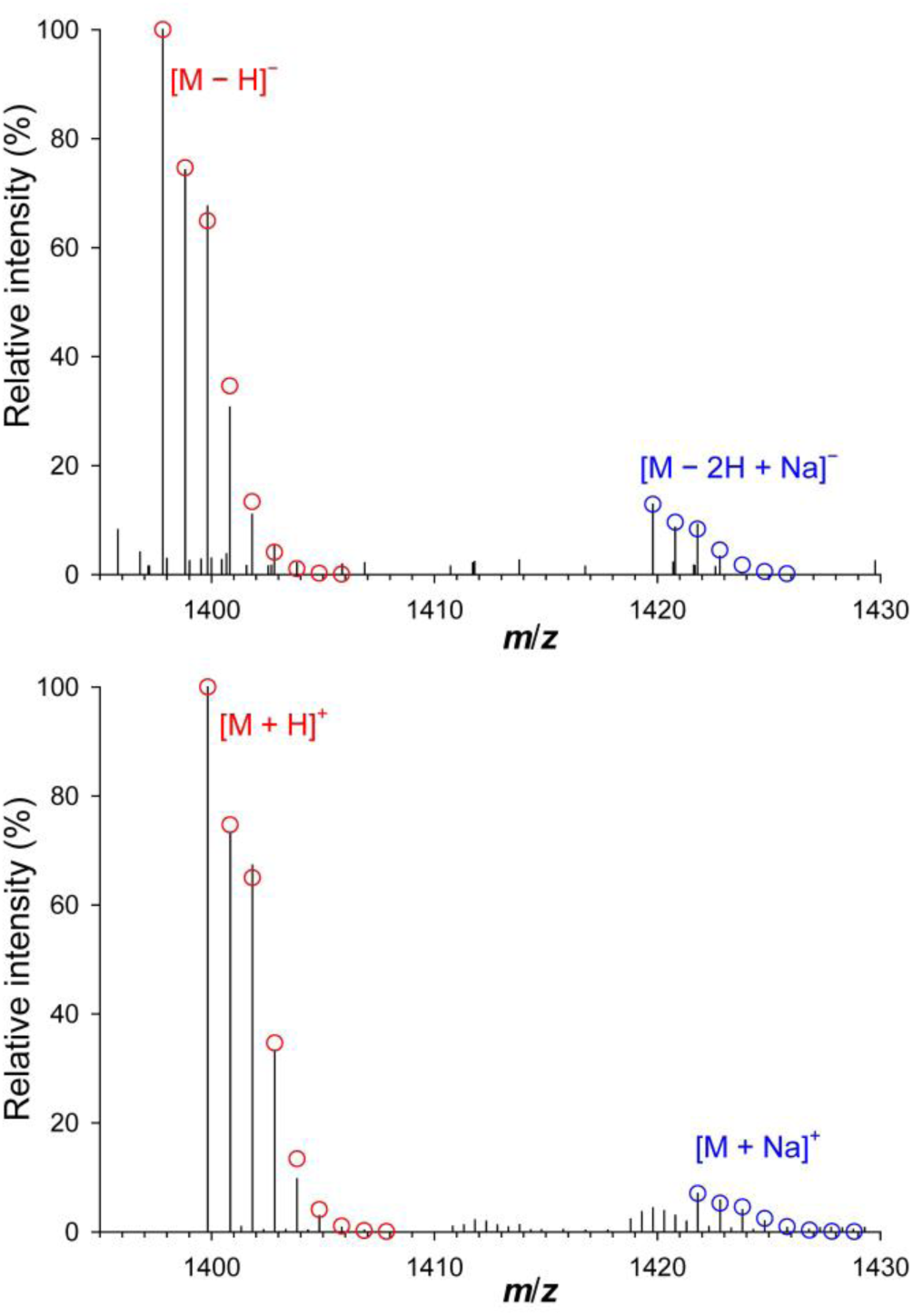
Analysis of the relative isotopologue abundances of the prominent adduct ions in LC–HRMS in positive and negative ionization modes ([M + H]^+^, [M + Na]^+^, [M − H]^−^, and [M − 2H + Na]^−^) of leadbeaterin-1(**1**) from fraction D7, establishing that leadbeaterin-1(**1**) has the elemental composition C_67_H_127_ClO_27_. The black lines show the LC– HRMS chromatogram, and the circles show the calculated relative isotopologue abundances for each adduct ion cluster.

More detailed analysis of the LC–MS data from fraction D7 revealed the presence of several minor metabolites eluting close to leadbeaterin-1(**1**), exhibiting similar *m*/*z* but with varied chlorination patterns. The characteristic isotopic pattern of chlorine helped to determine the degree of chlorination of individual molecular features. In addition to leadbeaterin-1(**1**), at least five additional analogues (**2** – **6**) were detected (**Table 3**). The molecular formulas were predicted based on the accurate masses of their adduct ions, isotopologue patterns, and comparisons of MS/MS spectra with those of leadbeaterin-1 (**1**), altogether supporting their structural similarity. Among others, the formula for a compound with [M + H]^+^ at *m*/*z* 1397.8109 (**2**) showed it to probably contain an extra double bond relative to leadbeaterin-1 (**1**), resulting in a slightly shorter retention time for compound **2** on the reversed-phase column (**Figure 8**).

**Figure 8.**
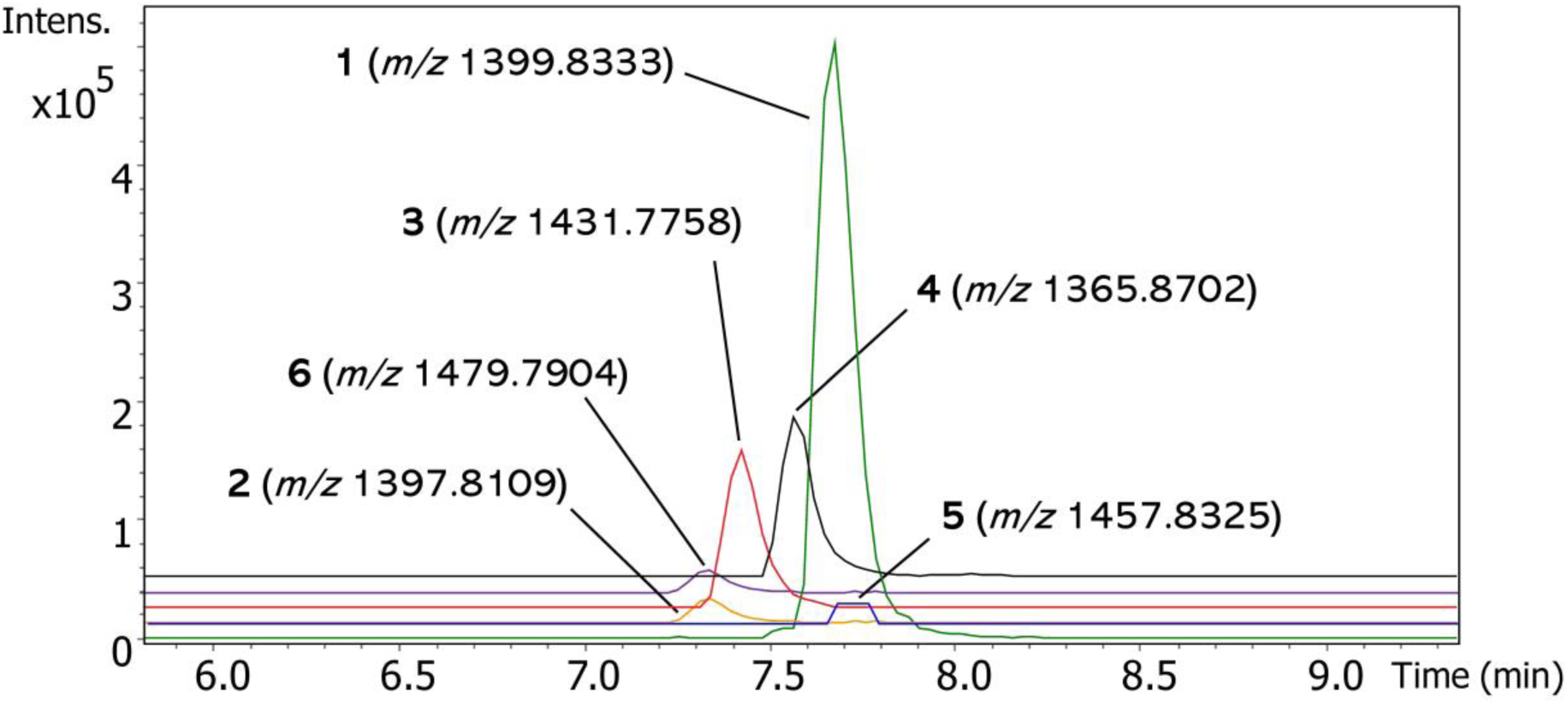
Extracted-ion chromatograms (EIC) of [M+H]^+^ of leadbeaterins (**1-6**) from the MeOH extract of UIO 394. The extraction windows are for the exact *m*/*z* ±0.02.

**Table 3.** Overview of the leadbeaterin analogues detected in *C. leadbeateri*.

| Proposed name | Accurate $m/z$ $[M+H]^+$ | Assigned formula* | No. Cl | $\Delta$ (ppm) | DBE |
| --- | --- | --- | --- | --- | --- |
| Leadbeaterin-1 (**1**) | 1399.8333 | $C_{67}H_{127}ClO_{27}$ | 1 | 1.5 | 4 |
| **2** | 1397.8109 | $C_{67}H_{125}ClO_{27}$ | 1 | 4.3 | 5 |
| **3** | 1431.7758 | $C_{67}H_{124}Cl_2O_{27}$ | 2 | 1.5 | 5 |
| **4** | 1365.8702 | $C_{67}H_{128}O_{27}$ | 0 | 1.0 | 4 |
| **5** | 1457.8325 | $C_{69}H_{129}ClO_{29}$ | 1 | 3.8 | 5 |
| **6** | 1479.7904 | $C_{67}H_{127}ClO_{30}S$ | 1 | -0.7 | 4 |

\*Neutral formula

Likewise, a compound with [M + H]^+^ at *m*/*z* 1431.7758 (**3**) appeared to be a variant of leadbeaterin-1 (**1**) containing an additional chlorine atom in addition to an extra double bond. The compound with [M + H]^+^ at *m*/*z* 1365.8702 (**4**) include without chlorine and the compound with [M + H]^+^ at *m*/*z* 1457.8325 (**5**) with the increased number of carbon atoms compared to leadbeaterin-1 (**1**) were also identified. Furthermore, the compound with [M + H]^+^ at *m*/*z* 1479.7904 (**6**) was determined to be C_67_H_127_ClO_30_S, differing from leadbeaterin-1 by the mass of an SO_3_ group, which suggested the presence of a sulfate group. The presence of a sulfate is supported by the peak’s relatively high intensity in negative ionization mode (data not shown) and the earlier elution of compound **6** from the reversed-phase column relative to leadbeaterin-1 (**1**), along with a prominent neutral loss of SO_3_ in its positive MS/MS spectrum to give a product-ion at *m*/*z* 1399.8303. Notably, all three strains of *C. leadbeateri* produced all six leadbeaterin analogues.

Given the molecular formulas and the observed bioactivity associated with fractions containing the leadbeaterins, we hypothesize they are likely structurally similar to another well-known family of toxins, namely the karlotoxins. These compounds are well-established as fish-killing toxins and exhibit a diverse array of structural derivatives (Van Wagoner et al., 2008; Van Wagoner et al., 2010). Upon comparing several karlotoxins with leadbeaterins, we found similarities in mass range, RDBEs, and patterns of chlorination and sulfation (**Table S2**). Karlotoxins typically feature a terminal diene, often chlorinated, contributing to their distinctive UV spectra, in contrast to the conjugated trienes commonly found in amphidinols (Krock et al., 2017). In our study, the observed maximum UV absorption for leadbeaterin-1 at 218 nm indicated the absence of a conjugated diene structure in the leadbeaterins (**Figure S4**), diverging from typical karlotoxins.

### 3.4. *In situ* detection of leadbeaterins from the fish killing event

Having access to samples collected during the bloom in 2019, we next set out to investigate whether we could detect leadbeaterins retrospectively in extracts from bloom material associated with the fish kills in 2019. The bloom was initially observed within fjord systems in Nordland county and later extended to Troms county (John et al., 2022). We detected leadbeaterins in the water sample from the Troms algal bloom. The samples were directly extracted from the bloom water and applied to a C_18_ SPE cartridge to recover the toxins. Analysis of the LC–MS data confirmed the presence of leadbeaterins (**Figure 9**).

**Figure 9.**
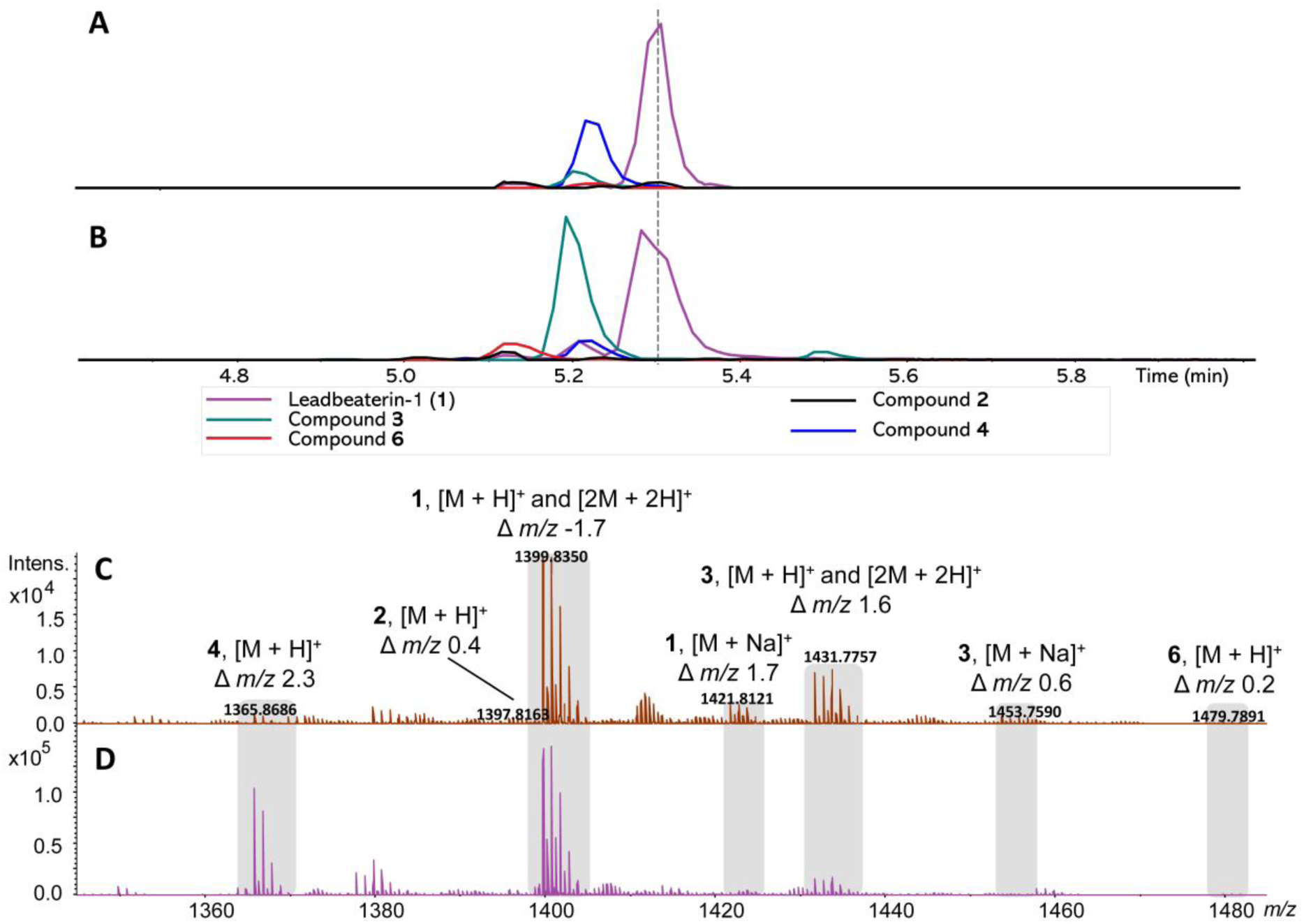
LC–HRMS EICs of [M+H]^+^ and the LC–HRMS spectra of leadbeaterins. The extraction window is for the exact *m*/*z* ±0.02. **A**. EICs of MeOH extracts generated after SPE from bloom water sample. **B**. EICs of MeOH extract of cells from UIO 394. **C**. LC–HRMS spectrum from water sample. **D**. LC–HRMS spectrum from MeOH extract of cells from UIO 394. The HRMS-spectrum is shown for the retention time window of 5.18–5.34 min.

## 4. Discussion

The presence of *C. leadbeateri* in the Northern Norwegian fjords during the 2019 event, leading to fish kills, has been confirmed by electron microscopical examination and through metabarcoding DNA-analysis of seawater samples, and barcoding of the strains from the 2019 event (John et al., 2022). However, the precise mechanism of fish mortality and the nature of the ichthyotoxins produced by *C. leadbeateri* have remained elusive. Our primary objective was to investigate and identify the fish-killing toxins within *C. leadbeateri* strains.

The utility of the RTgill-W1 gill cell line for toxin identification has been well-documented in previous studies, effectively identifying numerous fish-killing toxins. For example, previous studies established that RTgill-W1 cells are sensitive to two brevetoxins, BTX-2 and BTX-3 isolated from *Karenia brevis*, as well as the karlotoxin KmTx-2 from *Karlodinium veneficum* and saxitoxins from *Alexandrium* spp. (Place et al., 2012; Dorantes-Aranda et al., 2015; Dorantes-Aranda, 2023). These findings have all validated the efficacy of this cell line for correlating *in vitro* toxicity with *in vivo* effects. In line with these findings, we successfully employed a bioassay-guided fractionation approach using RTgill-W1, allowing us to pinpoint putative ichthyotoxic compounds. This study marks the first successful identification of potential fish-killing toxins in *C. leadbeateri*, laying the foundation for subsequent studies focused on isolation of the individual toxins for structure elucidation and comparison of their individual toxicities and underlying modes-of-action.

Our investigation involved three strains of *C. leadbeateri* isolated from different years and locations in Northern Norway, with the primary aim of determining whether a common class of toxins was responsible for the fish mortality events observed in both the 1991 and 2019 blooms. Initial findings revealed a remarkable similarity in the chemical profiles among various *C. leadbeateri* strains associated with these events, strongly indicating that the blooms in both 1991 and 2019 were likely caused by the same ichthyotoxins, supporting our hypothesis for a common toxin class produced by different strains of *C. leadbeateri*. Additionally, the molecular networking approach (Wang et al., 2016) proved particularly effective in elucidating the primary classes of molecules produced by these strains. This approach highlighted the complexity and variety in their metabolic profiles. The identification of compounds such as fatty acid amides, fatty acids, phospholipids, glycerolipids, and other lipids underscores a rich lipid metabolism within these *C. leadbeateri* strains.

By using the RTgill-W1 bioassay, we observed the consistent presence of highly toxic compounds within specific bioactive fractions across different strains from both the 1991 and 2019 blooms. This observation implies commonalities in toxin production among these strains. The application of PCA facilitated the visualization of relationships between these fractions. By clustering fractions with similar compound profiles, we were able to identify unique characteristics among the outliers. Furthermore, heatmap analysis highlighting several compounds exhibiting significantly high abundance within bioactive fractions. PCA and heatmaps provided a powerful tool to pinpoint potential candidates responsible for toxicity.

One of the key outcomes of our study is the identification of a specific compound, which we have named leadbeaterin-1 ([M + H]^+^ of *m*/*z* 1399.8333), as a potential toxin. This compound, with an elemental composition of C_67_H_127_ClO_27_, represents a notable discovery in the context of fish-killing toxins. The consistently high bioactivity associated with fractions containing this compound across each strain highlights its potential as a major contributor to the toxicity of the investigated algal strains. Moreover, our results emphasize the critical nature of fish toxin discovery by demonstrating that identical algal toxins can be responsible for multiple fish mortality events over different times and locations.

Additionally, the discovery of leadbeaterin analogues with variations in double bonds, chlorine and sulfate content, and number of carbon atoms highlights the structural diversity inherent in these compounds. These findings strongly suggest that the leadbeaterins produced by these algal strains are not restricted to a single compound. Instead, they likely represent a spectrum of structurally related compounds, each with potentially varying degrees of toxicity.

Our detailed MS analysis revealed that the molecular formulas of the leadbeaterins from *C*. *leadbeateri* are strikingly similar to those of the known karlotoxins, which is why we hypothesize that the leadbeaterin toxins are also long linear polyhydroxylated polyketides. The karlotoxin family stands out as a unique and well-studied group of fish-killing toxins. Their modes of action have been proposed (Waters et al., 2015), providing a foundation for understanding the putative toxins we identified in this study. This apparent connection with the karlotoxin family not only expands our understanding of leadbeaterins, but also provides a valuable reference point for further investigations into their biological activities. More intriguingly, this discovery further supports an evolutionary connection between haptophytes and dinoflagellates in the family Kareniaceae (Tengs et al., 2000). Members in this family possess a chloroplast type that is assumed to originate from the tertiary endosymbiosis of a haptophyte (Gabrielsen et al., 2011; Dorrell and Howe, 2012). The presence of karlotoxin-like compounds in the haptophyte *C. leadbeateri* raises the possibility of an ancestral link in the evolution of the genera *Karlodinium* and *Chrysochromulina.* To fully unravel this evolutionary narrative, there is a need for in-depth genomic, biochemical and phylogenetic investigations between these genera of microalgae.

The *in situ* detection of the putative toxin analogues within a natural algal bloom is of paramount importance. It indicates that these compounds are not confined to the controlled conditions of a laboratory. Rather, our investigations strongly indicate that they were also present in the ecosystems where fish mortality events were occurring, emphasizing their potential real-world relevance. Additionally, this *in situ* detection highlights the urgent need for continuous monitoring and comprehensive understanding of the production of these ichthyotoxins within their natural environments, and even if leadbeaterins turn out not to be ichthyotoxic themselves, they could make excellent biomarkers for the presence of ichthyotoxic *C. leadbeateri* blooms.

## 5. Conclusion

In conclusion, our combined approach of bioassay-guided isolation and advanced analytical techniques enabled us to make significant progress in the identification and initial characterization of the likely causative ichthyotoxic compounds produced during not only the most recent *C. leadbeateri* harmful algal blooms in Northern Norwegian fjords in 2019, but likely also of the 1991 bloom. The discovery of these putative fish-killing toxins, their structural diversity, and their potential links to known toxin families open up exciting possibilities for further research. These include further purification and structure elucidation of the toxins, as well as a deeper understanding of their modes of action and factors affecting toxin production. Importantly, this research holds significant potential for practical applications, particularly in developing robust monitoring and mitigation strategies to address the ecological and economic impacts of these devastating fish-killing events.

## CRediT authorship contribution statement

**Xinhui Wang:** Conceptualization, Methodology, Validation, Investigation, Writing – original draft, Visualization, Project administration. **Mathias Fon:** Conceptualization, Methodology, Validation, Investigation, Writing – original draft, Visualization. **Aaron J.C. Andersen:** Methodology, Resources, Investigation. **Anita Solhaug:** Methodology, Investigation, Writing – original draft. **Richard A. Ingebrigtsen:** Methodology, Resources, Writing – review & editing. **Ingunn A. Samdal:** Investigation, Funding acquisition, Writing – review & editing, Project administration. **Silvio Uhlig:** Investigation, Writing – review & editing, Funding acquisition, Project administration. **Christopher O. Miles:** Methodology, Investigation, Writing – review & editing. **Bente Edvardsen:** Investigation, Writing – review & editing, Project administration. **Thomas O. Larsen:** Investigation, Writing – review & editing, Visualization, Project administration.

## Declaration of competing interest

The authors declare that they have no known competing financial interests or personal relationships that could have appeared to influence the work reported in this paper.

## Acknowledgments

This work was supported by the Norwegian Research Council for the project Toxic microalgae in Norwegian waters − Uncovering fish-killing mechanisms of phytoplankton from Scandinavian waters (ToxANoWa) (project no. 314861/E40). The authors acknowledge Xu Chen, Kah Yean Lum and Manca Vertot for their contributions to algal cultivation and helpful discussions, as well as Wenche Eikrem and Luka Šupraha for access to the *C. leadbeateri* strains.

## Notes

### Competing Interest Statement

The authors have declared no competing interest.

