## Supplementary information for "Insights into the nature of the microalgal toxins from the *Chrysochromulina leadbeateri* blooms in Northern Norwegian fjords"

^*^Shared first authorships

**Table of contents**

**Figure S1.** Base peak chromatograms (BPCs) of the three *C. leadbeateri* strains (UIO 035, UIO 393 and UIO 394) in positive ionization mode. Identities of selected peaks are indicated.

**Figure S2.** Base peak chromatograms (BPCs) of fraction D in positive ionization mode across each strain.

**Figure S3.** Base peak chromatograms (BPCs) of 15 subfractions from fraction D combined from the UIO 394 and UIO 035 strains in the positive ionization mode.

**Figure S4.** The LC–UV spectrum of *m*/*z* 1399 (leadbeaterin-1), showing absorptions at 196 nm and 218 nm.

**Table S1.**  Viability of the RTgill-W1 cell line after exposure to fractions A–E from the MeOH extracts *of C. leadbeateri* strains.

**Table S2.** Overview of selected Karlotoxins

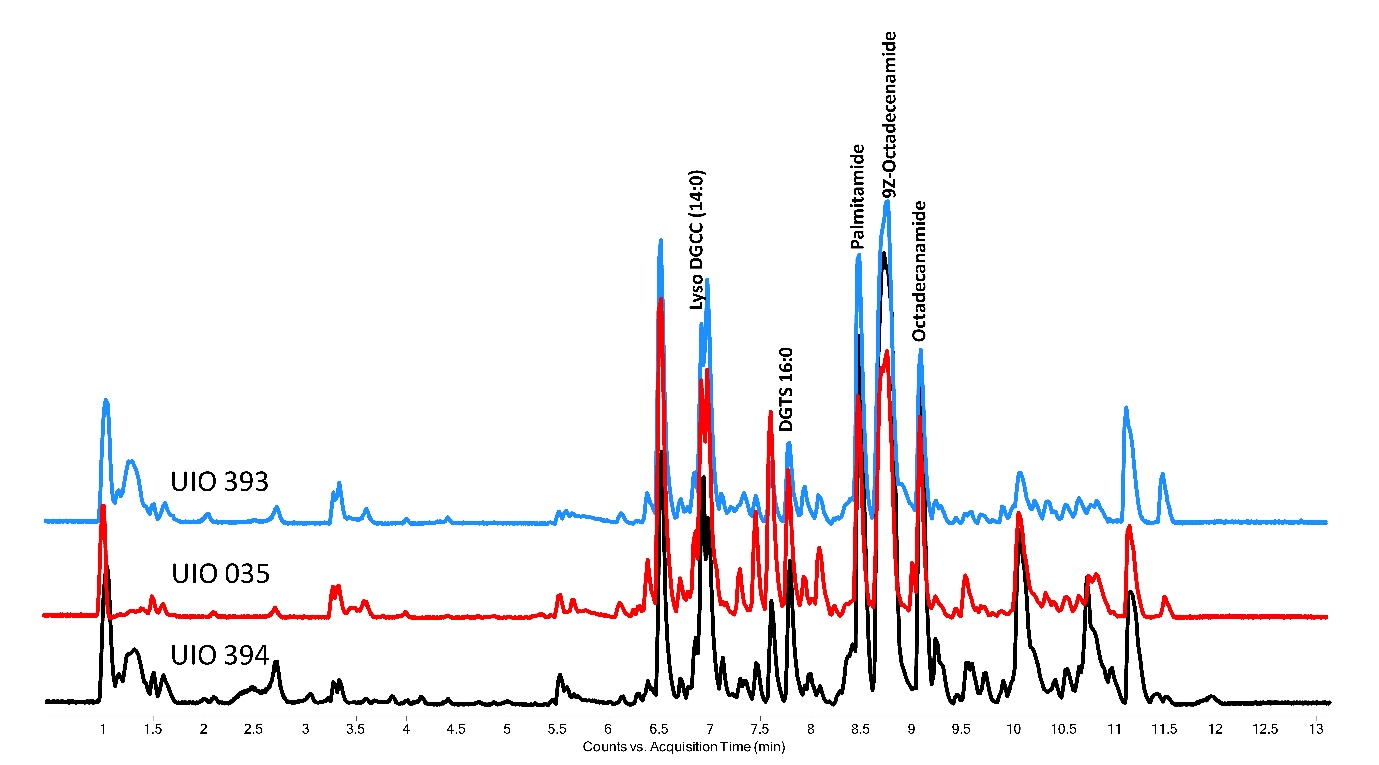

**Figure S1.** Base peak chromatograms (BPCs) of the three *C. leadbeateri* strains (UIO 035, UIO 393 and UIO 394) in positive ionization mode. Identities of selected peaks are indicated.

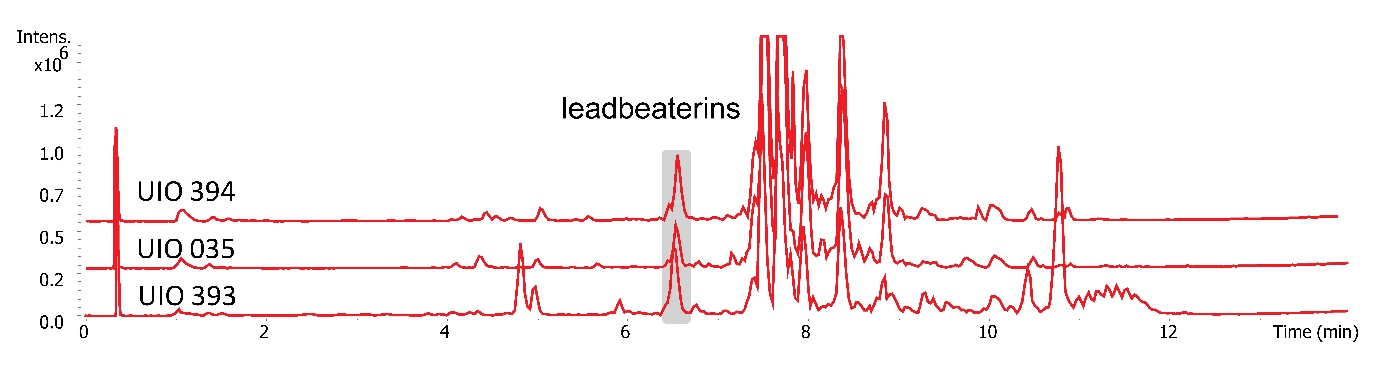

**Figure S2.** Base peak chromatograms (BPCs) of fraction D in positive ionization mode across each strain.

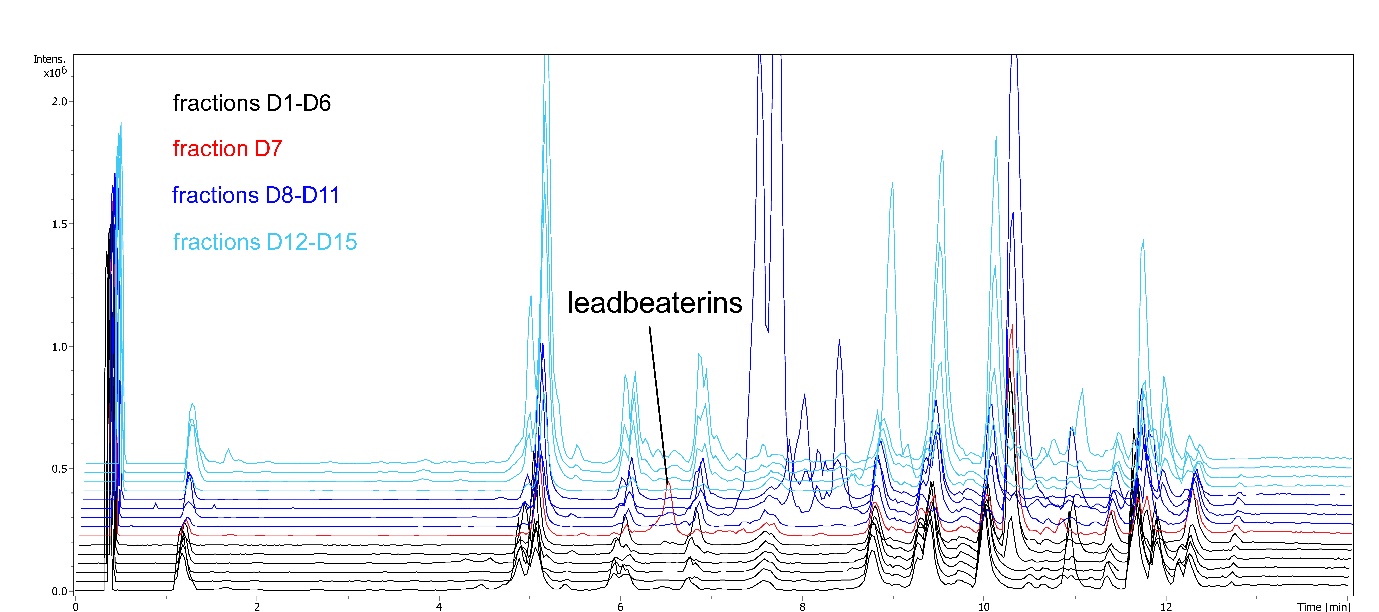

**Figure S3.** Base peak chromatograms (BPCs) of 15 subfractions from fraction D combined from the UIO 394 and UIO 035 strains in the positive ionization mode.

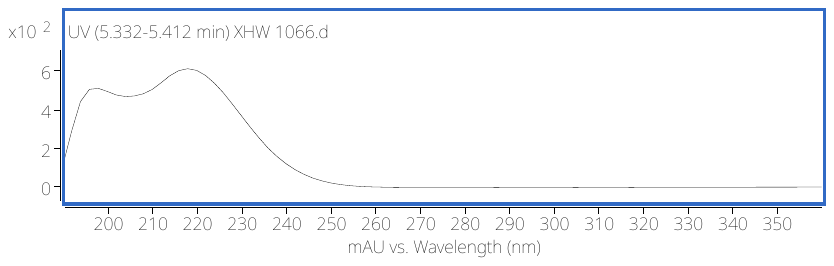

**Figure S4.** The LC–UV spectrum of *m*/*z* 1399 (leadbeaterin-1), showing absorption maxima at 196 nm and 218 nm.

**Table S1.** Viability of the RTgill-W1 cell line after exposure to fractions A–E from the MeOH extracts *of C. leadbeateri* strains. The fractions were obtained by C18 flash chromatography.

| **Strains** | **Fractions A–E** | **Concentration (1%, per well, mg/µL)** | **Cellular Viability (% of control)** |
| --- | --- | --- | --- |
| UIO 394 | **A.** 20–40% MeOH | 0.00275 | 33 |
|  | **B.** 50–60% MeOH | 0.0004 | 98 |
|  | **C.** 70–80% MeOH | 0.0009 | 100 |
|  | **D.** 90% MeOH | 0.0003 | 0.61 |
|  | **E** 100% MeOH | 0.0035 | 0.69 |
| UIO 393 | **A.** 20-40% MeOH | 0.00012 | 110 |
|  | **B.** 50-60% MeOH | 0.00014 | 110 |
|  | **C.** 70-80% MeOH | 0.0001 | 120 |
|  | **D.** 90% MeOH | 0.00009 | 1.0 |
|  | **E.** 100% MeOH | 0.00013 | 1.2 |
| UIO 035 | **A.** 20-40% MeOH | 0.0001 | 110 |
|  | **B.** 50-60% MeOH | 0.00012 | 110 |
|  | **C.** 70-80% MeOH | 0.00002 | 120 |
|  | **D.** 90% MeOH | 0.00008 | 0.63 |
|  | **E.** 100% MeOH | 0.00028 | 0.94 |

| **Name** | **Molecular Formula** | **Monoisotopic Mass** | **Source** | **UV maximum** | **References** |
| --- | --- | --- | --- | --- | --- |
| Karlotoxin 1 | C_69_H_126_O_24_ | 1338.8639 | *Karlodinium veneficum* | 225 | (Van Wagoner et al., 2008) |
| Karlotoxin 2 | C_67_H_121_ClO_24_ | 1344.7936 | *Karlodinium veneficum* | 235 | (Peng et al., 2010) |
| Karlotoxin 3 | C_68_H_124_O_24_ | 1324.8483 | *Karlodinium veneficum* | 228 | (Van Wagoner et al., 2010) |
| 10-*O*-sulfo-karlotoxin 3 | C_68_H_124_O_27_S | 1404.8050 | *Karlodinium veneficum* | 228 | (Van Wagoner et al., 2010) |
| 65*E*-chloro-karlotoxin 1 | C_69_H_125_ClO_24_ | 1372.8249 | *Karlodinium veneficum* | 233 | (Van Wagoner et al., 2010) |

**Table S2.** Overview of selected Karlotoxins.
